# Genome-resolved surveillance of African *Klebsiella oxytoca* species complex genomes reveals resistome-mobilome and biosynthetic gene cluster diversity

**DOI:** 10.64898/2026.08.25.747093

**Authors:** Samweli Bahati, Abdalah Makaranga, Lesley Hoyles, Reuben S. Maghembe

## Abstract

The *Klebsiella oxytoca* species complex (KoSC) comprises taxonomically diverse commensals and opportunistic pathogens, but its genomic diversity remains poorly characterized across Africa. We curated publicly available African KoSC data through raw-read and public-assembly routes and analyzed 163 African genomes together with 282 global comparators. Pangenome, phylogenomic, sequence-typing, surface-locus, antimicrobial-resistance, plasmid-replicon, mobile-element, biosynthetic-gene-cluster, and virulence-component analyses were integrated. The African collection comprised *K. michiganensis* (112/163), *K. oxytoca* (36/163), *K. pasteurii* (8/163), and *K. grimontii* (7/163) from 13 countries. The African pangenome contained 4,286 core, 4,505 shell, and 18,066 cloud gene families. Official PubMLST sequence types were assigned to 129/163 genomes. Four core/intrinsic antimicrobial-resistance-associated loci (*ompA, oqxA, oqxB,* and *bla*OXY) occurred in all genomes, whereas acquired resistance determinants were heterogeneous. Intact *til* biosynthetic gene clusters occurred in 55/163 genomes and intact *leup* clusters in 103/163. *leup* was concentrated in *K. michiganensis* (102/112), whereas intact *til* was frequent in *K. oxytoca* (26/36), *K. pasteurii* (7/8), and *K. grimontii* (7/7). *Klebsiella*-focused Virulence Factor Database screening detected at least one curated component in 112/163 genomes, but no complete curated factor; three *K. michiganensis* genomes carried complete *mrkABCDF* structural-operon candidates. These data define an African genome-resolved baseline for KoSC diversity and identify species-structured biosynthetic loci alongside heterogeneous resistance and mobilome profiles.

**IMPORTANCE:** The *Klebsiella oxytoca* species complex contains opportunistic pathogens that can carry antimicrobial-resistance genes and biosynthetic pathways linked to host-associated effects, yet African representatives remain poorly characterized. This study provides a curated genomic view of the complex across 13 African countries and places those genomes in a global context. The analysis shows extensive variation in accessory genes, resistance determinants, and mobile genetic elements, while two biologically important biosynthetic loci have strikingly different species distributions. These findings provide a reproducible baseline for genomic surveillance of this understudied *Klebsiella* group in Africa and identify specific lineages and genomic features for future population-based sampling, complete-genome reconstruction, and laboratory validation.

## INTRODUCTION

The *Klebsiella oxytoca* species complex (KoSC) comprises 11 phylogroups that include *K. michiganensis* (Ko1 and Ko5), *K. oxytoca* (Ko2), *K. spallanzanii* (Ko3), *K. pasteurii* (Ko4), *K. grimontii* (Ko6), *K. huaxiensis* (Ko8) and “*K. mammaliorum*” (Ko12), together with three unnamed taxa (Ko7, Ko10 and Ko11) (1, 2). KoSC members occupy human, animal, and environmental reservoirs and include gastrointestinal commensals as well as opportunistic pathogens associated with urinary tract infections, bacteraemia, healthcare-associated outbreaks, and antibiotic-associated haemorrhagic colitis (3). This taxonomic diversity is poorly resolved by conventional phenotypic identification: genome-wide analyses are required for reliable species assignment and have revealed substantial misclassification among publicly deposited *Klebsiella* genomes (3, 4).

Genome-based studies have further shown that KoSC diversity extends beyond species boundaries to population structure, surface-antigen loci and antimicrobial resistance (AMR) determinants. Analyses of public genomes have identified extensive sequence-type and K-and O-locus diversity, the core chromosomal *bla*_OXY_ lineage, and heterogeneous repertoires of acquired resistance and virulence-associated genes (3, 4). High-resolution genomic typing is consequently central to KoSC surveillance because resistance determinants are distributed across genetically diverse lineages and may occur within mobile genetic contexts that are not captured by species designation alone. African genomic-surveillance capacity has expanded substantially, but sequence availability remains uneven among countries and bacterial taxa, and recent genomic studies of *Klebsiella* from African clinical settings continue to expose gaps in taxonomic resolution, sampling breadth, and genomic representation (3, 5–7).

KoSC genomes also encode specialized-metabolite pathways with experimentally established host-associated activities. The *til* (tilimycin, also known as kleboxymycin) biosynthetic gene cluster (BGC) occurs in *K. oxytoca*, *K. pasteurii*, *K. grimontii,* and *K. michiganensis* and encodes the pathway responsible for tilimycin and tilivalline biosynthesis (1, 8). Tilimycin is genotoxic, whereas tilivalline targets microtubules, providing distinct molecular mechanisms underlying the cytotoxic activity associated with the locus (9). A second BGC, *leup*, encodes leupeptin biosynthesis and was initially found to be enriched among clinical respiratory isolates belonging to *K. michiganensis* (10). Subsequent work demonstrated that *N*-acetylneuraminic acid activates *leup* transcription and leupeptin-derived pyrazinone biosynthesis in *K. michiganensis*, linking this pathway to host-derived nutrient sensing (11). Genome-scale analysis has since shown an almost exclusive association of the complete *leup* operon with *K. michiganensis*, in which it was detected in 90.6% of genomes (1). The distribution of these two chromosomally encoded virulence-associated BGCs therefore represents an additional dimension of KoSC genomic diversity that is biologically distinct from AMR and mobilome variation.

Here, we assembled and taxonomically curated publicly available African KoSC genomes and placed them within a global phylogenomic context. We resolved African KoSC species and lineage structure, pangenome diversity and surface-locus variation; characterized the distribution of AMR determinants, plasmid replicons and non-plasmid mobile genetic elements (MGEs); and determined the species distribution and genomic architecture of the *til* and *leup* BGCs. We additionally assessed Klebsiella-labelled virulence-associated components across the African collection. These analyses provide a genome-resolved baseline for KoSC diversity and surveillance in Africa.

## MATERIALS AND METHODS

### Genome acquisition, assembly and taxonomic curation

Public African KoSC records were curated from NCBI sequence resources using KoSC species, complex and phylogroup search terms together with African geographic terms. Records were screened for isolate traceability, sampling origin, sequencing strategy, accession continuity and sequence availability. Two acquisition routes were retained. First, paired-end Illumina isolates with available raw reads underwent read-level quality control with fastp v0.24.0 (12), *de novo* assembly with SPAdes v4.2.0 (13) and assembly assessment with QUAST v5.3.0 (Gurevich et al., 2013). Second, public assembly-only additions selected from NCBI-curated assembly resources were evaluated using accession provenance, deposited taxonomy, and assembly-quality information. Complete accession and provenance fields are provided in Supplementary Table S1, fastp metrics are in Supplementary Table S2, and QUAST metrics in Supplementary Table S3.

QUAST metrics included total assembly length, contig count and N50. Values of approximately 5.0-6.8 Mb, ≤1,000 contigs and N50 ≥10 kb were used as review thresholds. FastANI v1.34 (14) was used for taxonomic confirmation against a curated KoSC reference panel (Fig. 1). Genomes were retained when the highest qualifying match was a KoSC reference with ANI ≥95.0%, alignment-fraction proxy ≥0.20, and a best-versus-second-hit ANI difference ≥0.20.

**Figure 1.**
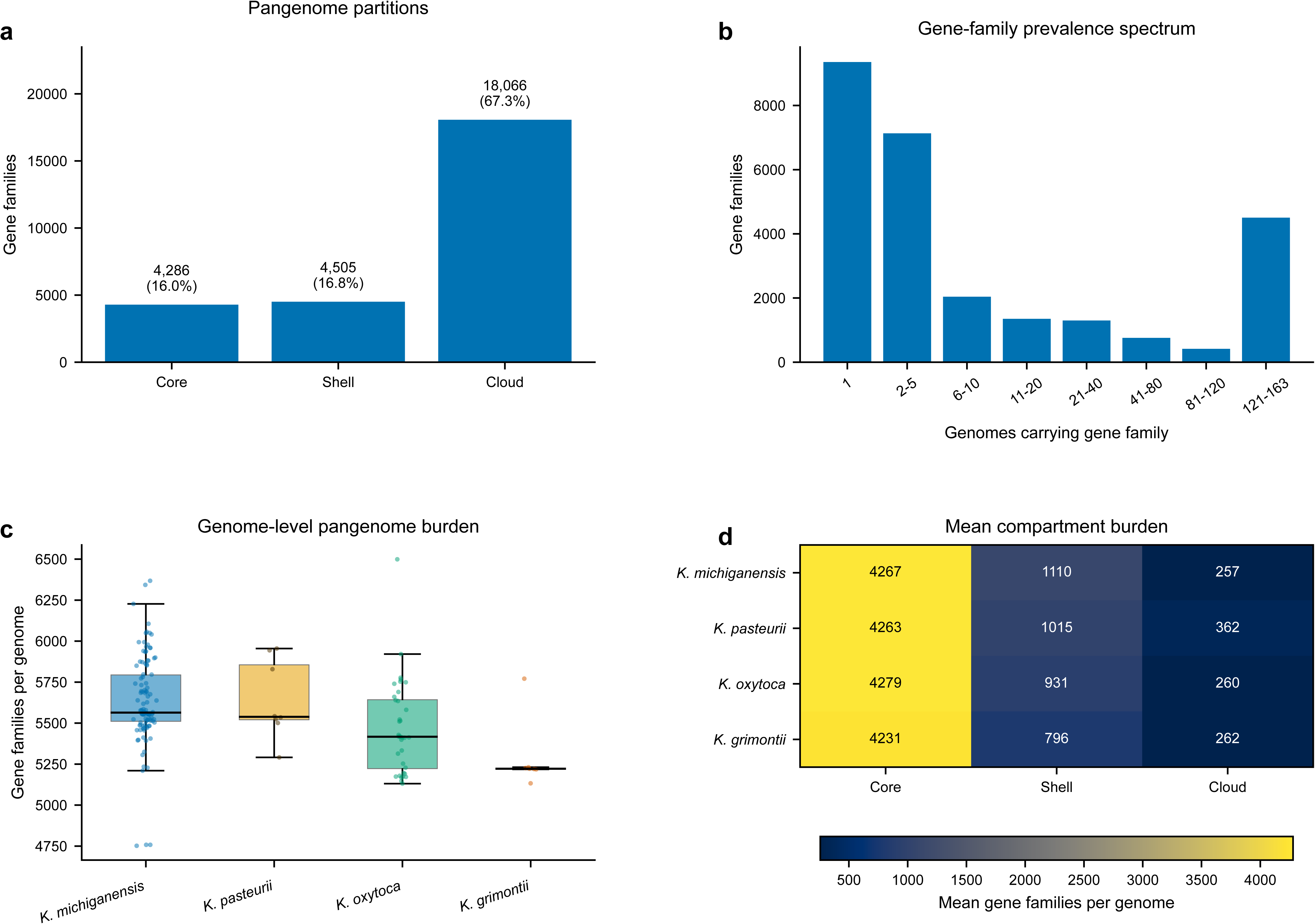
Construction of the African KoSC genome dataset and global comparator panel. Public African records were curated through raw-read and public-assembly routes. Paired-end Illumina data underwent fastp quality control, de novo assembly, QUAST review and taxonomic screening; public assembly additions were evaluated using accession provenance, assembly quality and curated taxonomy. The final downstream African set contained 163 genomes. A 282-genome global comparator panel produced the 445-genome combined context set. Detailed QC and taxonomy records are provided in Supplementary Tables S1–S4 and Supplementary Figure S1.

### Pangenome and phylogenomic analysis

Pangenomes were inferred with PPanGGOLiN v4.3.1 for the African primary set (n = 163) and the combined context set (n = 445) (15). PPanGGOLiN assigns gene families to persistent (i.e., core), shell, and cloud partitions. Partition counts, gene-family prevalence, and per-genome burdens were exported directly from the pangenome outputs. Persistent/core gene families were aligned and concatenated for maximum-likelihood phylogenetic inference with IQ-TREE v2.4.0 using ModelFinder model selection under the Bayesian information criterion (16).

### Sequence type and surface-locus assignment

KoSC multilocus sequence typing used the *K. oxytoca* PubMLST seven-locus scheme (*gapA, infB, mdh, pgi, phoE, rpoB* and *tonB*) and current curated profile definitions (17). Only official numeric PubMLST assignments are reported as sequence types (STs). Profiles without an official numeric assignment are reported as unassigned. Capsule (KL) and O-antigen (OL) loci were assigned with Kaptive v3.2.1 (18) using KoSC-specific locus databases (2).

### AMR and mobilome analyses

African assemblies were screened with ABRicate v1.4.0 against matched CARD, NCBI AMR, and ResFinder nucleotide database snapshots dated 10 July 2026. All three screens used ≥90% nucleotide identity and ≥60% reference coverage. CARD provides ontology-based curation of AMR determinants (19), and ResFinder provides a curated genotype framework for acquired resistance genes (20). The same assemblies were screened with AMRFinderPlus v4.2.7 using NCBI database version 2026-05-15.1 and default database scope (21). Standalone AMR features were added to the main binary matrix when accepted nucleotide evidence met ≥90% identity and ≥60% reference coverage. Database labels were harmonized at exact-key and conservative family levels while retaining source-specific labels; *qacE* and *qacE*Δ*1* were maintained as separate features. OqxAB was evaluated at the paired-locus level using same-contig, same-strand *oqxA*-*oqxB* evidence within 5 kb.

Plasmid replicons were detected with ABRicate/PlasmidFinder at ≥95% identity and ≥60% reference coverage (22). MOB-suite v3.1.9 mob_recon supplied plasmid-associated/non-chromosomal contig assignments (23). Non-plasmid mobile-element calls were retained after removal of ribosomal RNA features from the source matrix.

### Reference-guided *til* and *leup* BGC classification

Broad BGC screening used antiSMASH v7.1.0 (24). Final *til* and *leup* classifications were derived from reference-guided component detection and locus architecture. BLAST+ v2.17.0 supplied protein and targeted nucleotide similarity searches (25). For the *til* BGC, the reference was GenBank MF401554.1 and the 12 ordered components *mfsX, uvrX, hmoX, adsX, icmX, dhbX, aroX, npsA, thdA, npsB, npsC* and *marR*. Strict protein support required ≥90% amino-acid identity, ≥90% reference coverage, E-value ≤1×10−5 and a subject/reference length ratio of 0.75–1.30. Targeted nucleotide rescue used the same identity, coverage and E-value thresholds.

The *leup* reference was *K. oxytoca* chromosome NC_016612.1. Five operational components were evaluated: KOX_RS06945 (MFS transporter), KOX_RS06940 (LuxE/PaaK-family acyltransferase), KOX_RS06935 (acyl-CoA reductase), KOX_RS06930 (phenylacetate-CoA ligase-family protein) and KOX_RS06925 (GNAT-family N-acetyltransferase). Strict calls used the same ≥90% identity, ≥90% coverage, E-value, and length-ratio criteria, followed by nucleotide and architecture review. Intact *leup* status required recovery of all five components with supported order, strand, and colocalization; assembly-fragmented and no-qualifying-locus states were retained separately.

### Virulence-component screening

Virulence-associated components were screened with ABRicate v1.4.0 against the local Virulence Factor Database (VFDB) snapshot dated 10 July 2026 using ≥90% nucleotide identity and ≥90% reference coverage. Analysis was restricted to 128 *Klebsiella*-labelled VFDB reference entries representing 122 unique curated components across 14 factor groups (26). Factor-level completeness required recovery of every curated component assigned to a factor. Type 3 fimbrial architecture was additionally evaluated with a structural *mrkABCDF* rule requiring all five structural genes on one contig in consistent order and strand. Detected VFDB components, factor-level evidence, genome summaries and type 3 fimbrial candidates are provided in Supplementary Tables S13–S16.

### Descriptive analysis

The genome was the unit of analysis. Counts and percentages use explicit denominators, with n = 163 for Africa-focused downstream analyses, n = 282 for the global comparator panel, and n = 445 for combined-context analyses. Feature categories are non-mutually exclusive where genomes contain multiple genes, replicons, MGEs or VFDB components. No population-weighted prevalence estimates were calculated from the public genome collection.

## RESULTS

### African KoSC dataset and geographic representation

Sequential metadata curation, assembly-quality review and taxonomic confirmation resolved a defined African KoSC analysis dataset (Fig. 1; Supplementary Tables S1). Of 176 African candidate genomes entering FastANI confirmation, 164 met the predefined KoSC species-level criteria. One genome (KoSC109; SRA run SRR21101699; no deposited genome assembly accession) was subsequently removed after annotation/pangenome failure, yielding the African primary set used for prevalence-based analyses (n = 163). A separate panel of 282 confirmed global comparator genomes was retained for context, producing a 445-genome set for comparative pangenome and phylogenetic analyses.

The primary African dataset comprised 163 genomes from 13 countries (Fig. 1; Supplementary Table S1). *K. michiganensis* accounted for 112/163 genomes (68.7%), followed by *K. oxytoca* (36/163, 22.1%), *K. pasteurii* (8/163, 4.9%) and *K. grimontii* (7/163, 4.3%). Ethiopia contributed 65 genomes, South Africa 22, Egypt 17, Kenya 16 and Tanzania 12; the remaining 31 genomes were distributed across Namibia, Uganda, Nigeria, Malawi, Rwanda, Senegal, Tunisia and Mozambique. Ethiopia contained all four species represented in the final dataset. Species-by-country distributions are detailed in Supplementary Table S5.

### African pangenome structure

PPanGGOLiN partitioned the African pangenome into 4,286 core gene families (16.0%), 4,505 shell families (16.8%), and 18,066 cloud families (67.3%) (Fig. 2). Low-frequency families formed the largest part of the gene-family prevalence spectrum. Mean per-genome core-family counts were similar across species: 4,279 in *K. oxytoca*, 4,267 in *K. michiganensis*, 4,263 in *K. pasteurii,* and 4,231 in *K. grimontii*, while shell and cloud burdens varied among species.

**Figure 2.**
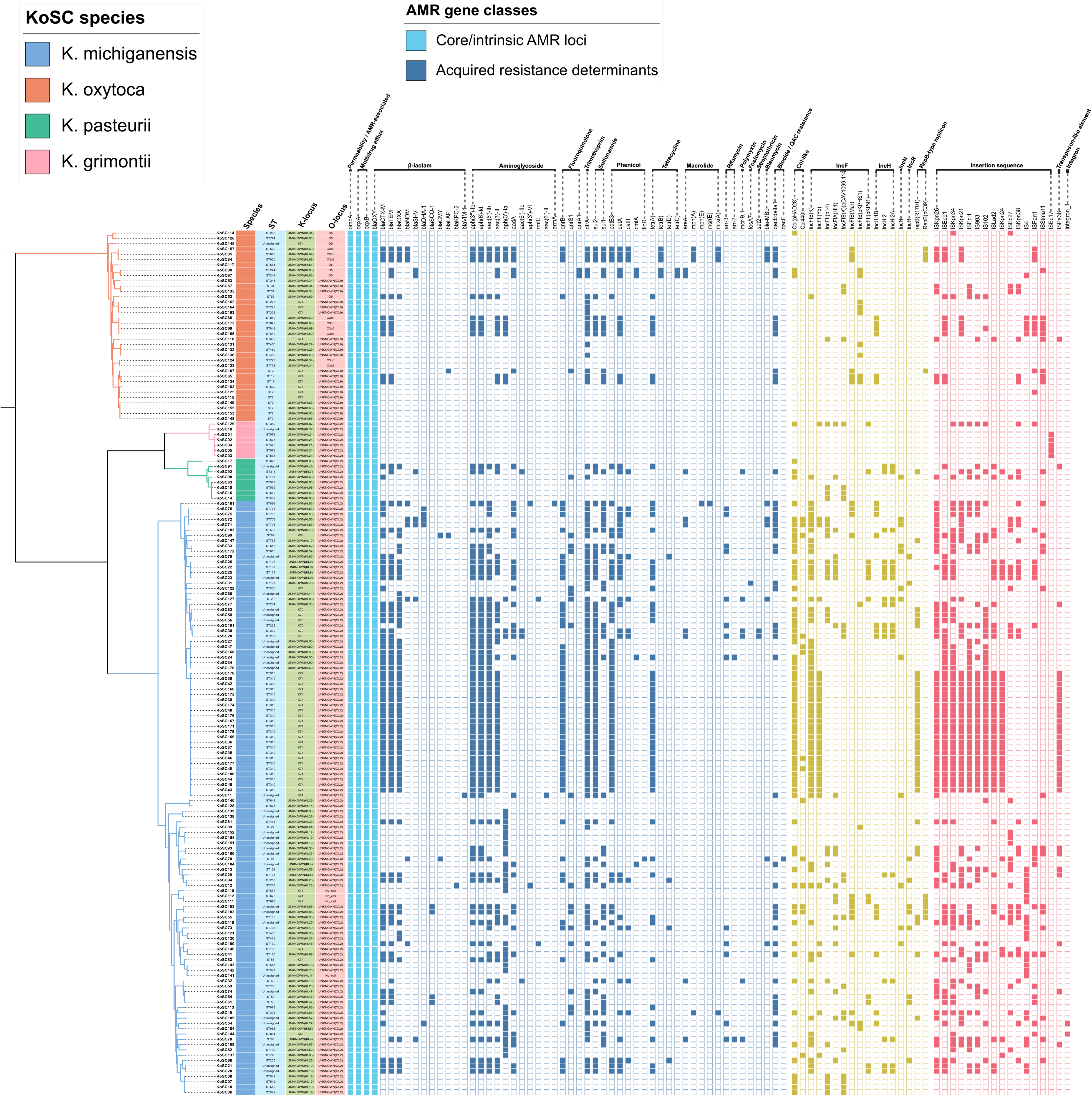
African KoSC pangenome structure. **a,** Distribution of 26,857 gene families among core (4,286; 16.0%), shell (4,505; 16.8%), and cloud (18,066; 67.3%) partitions in the 163-genome African set. **b,** Gene-family prevalence spectrum across the African genomes. **c,** Per-genome pangenome burden by species: K. michiganensis (n = 112), K. pasteurii (n = 8), K. oxytoca (n = 36) and K. grimontii (n = 7). **d,** Mean core, shell and cloud family counts by species. Core denotes the PPanGGOLiN persistent partition in the main-text presentation.

The combined 445-genome pangenome contained 4,116 core, 8,281 shell and 28,604 cloud gene families (Supplementary Figure S2), compared with 4,286, 4,505 and 18,066, respectively, in the African dataset. Mean persistent/core burden per genome was 4,137 in African genomes and 4,139 in global genomes; mean shell burden was 1,249 and 1,291 families, respectively.

### Phylogenomic structure and sequence typing

Persistent/core-gene phylogenies placed the African genomes across multiple species-associated branches within the global KoSC context (Fig. 3). Within the African dataset, *K. michiganensis* formed the largest species component, while *K. oxytoca*, *K. pasteurii* and *K. grimontii* occupied distinct species backgrounds. Official PubMLST assignments were available for 129/163 genomes (79.1%); 34 genomes (20.9%) remained unassigned. ST310 was the largest single sequence type (23 genomes, all *K. michiganensis*), followed by ST2 (7 *K. oxytoca*) and ST576 (5 *K. grimontii*) (Table 1).

**Figure 3.**
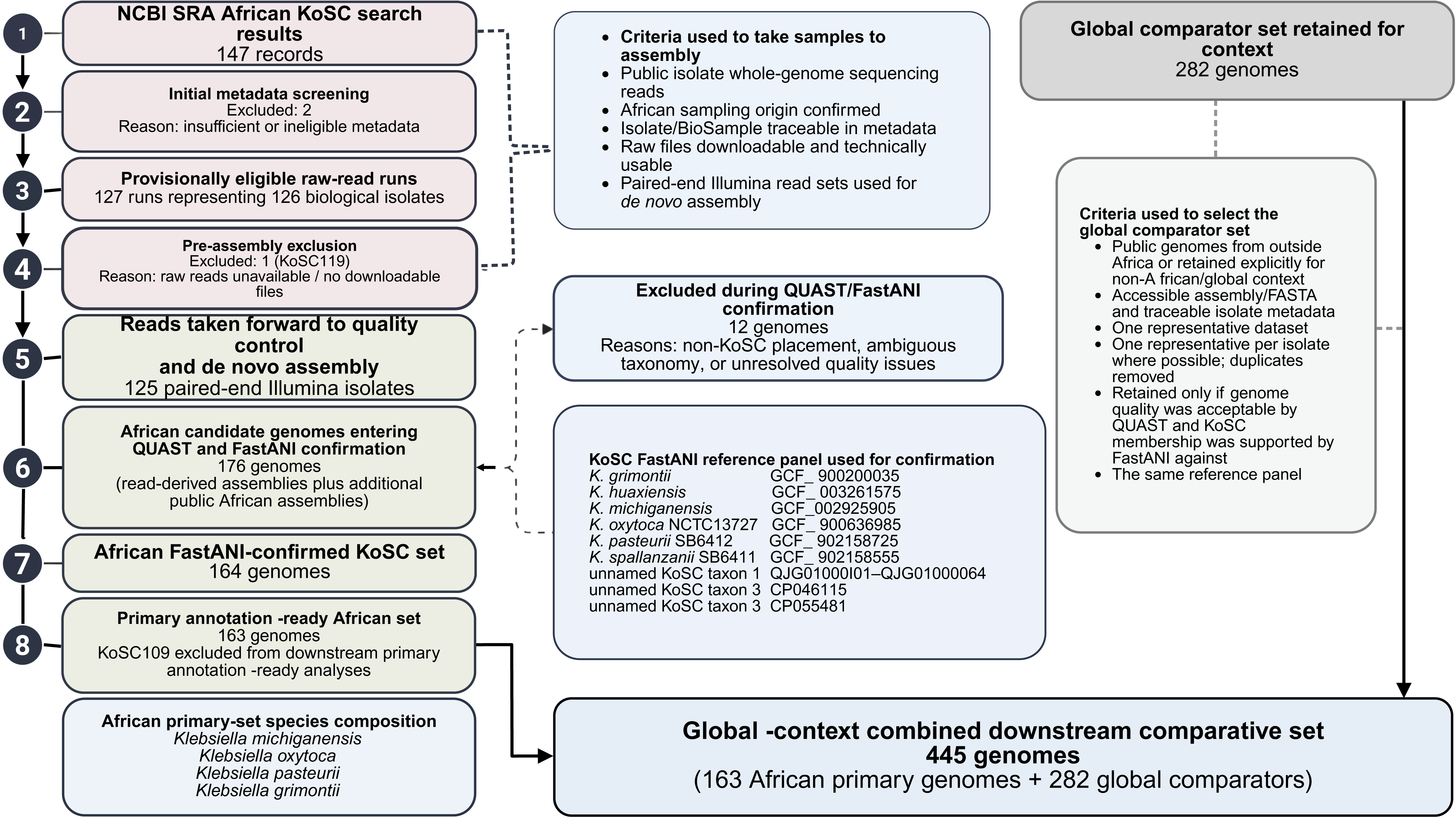
Persistent/core-gene phylogeny and til/leup BGC distribution in African KoSC genomes within global context. Maximum-likelihood trees were inferred independently from concatenated persistent/core-gene alignments. a, Combined 445-genome context phylogeny annotated by dataset, species, and current official sequence type. b, African 163-genome phylogeny annotated by species, country, current official sequence type, and intact til/leup BGC status. Profiles without an official PubMLST numeric assignment are labelled Unassigned. Both trees were midpoint-rooted for display.

**Table 1.**
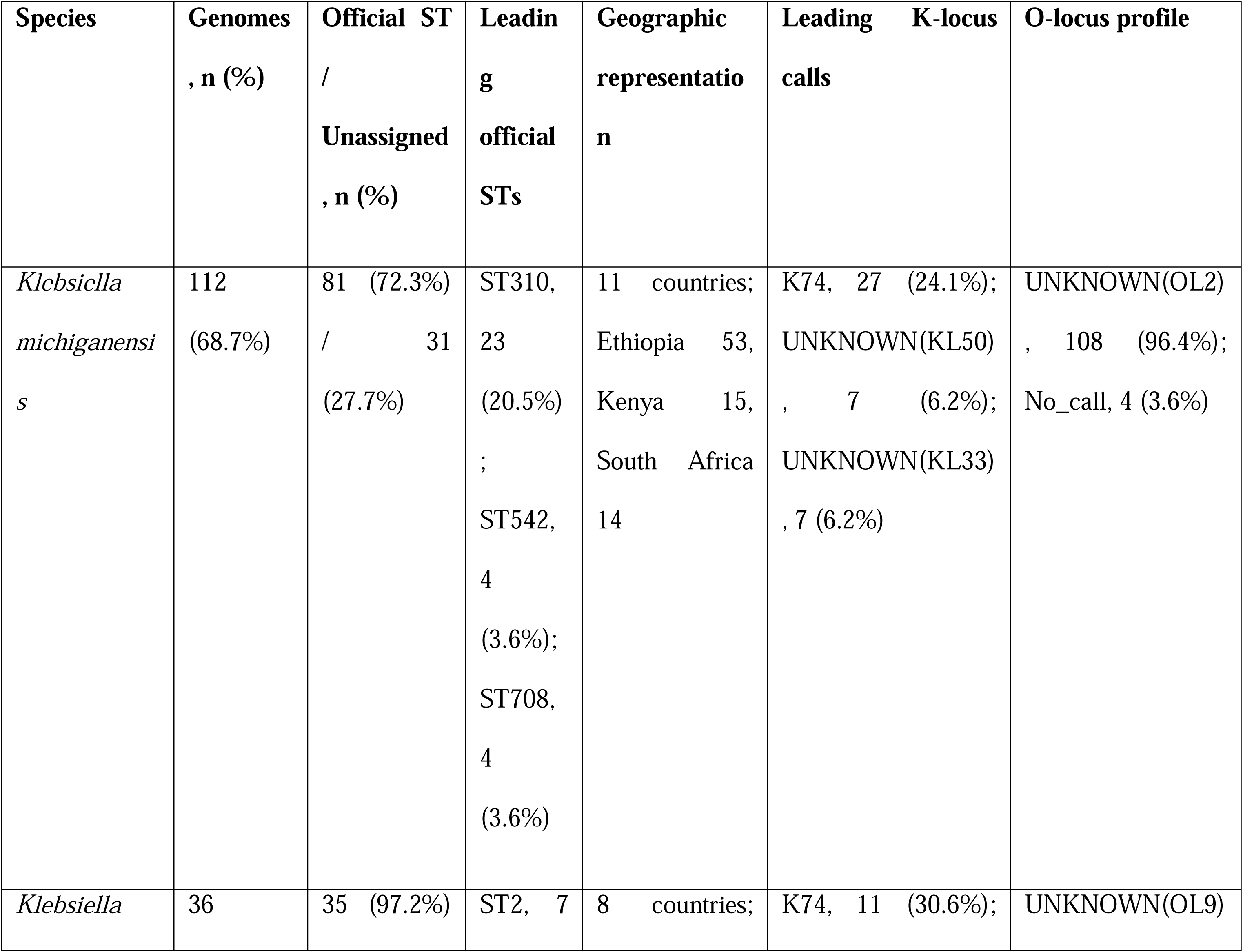
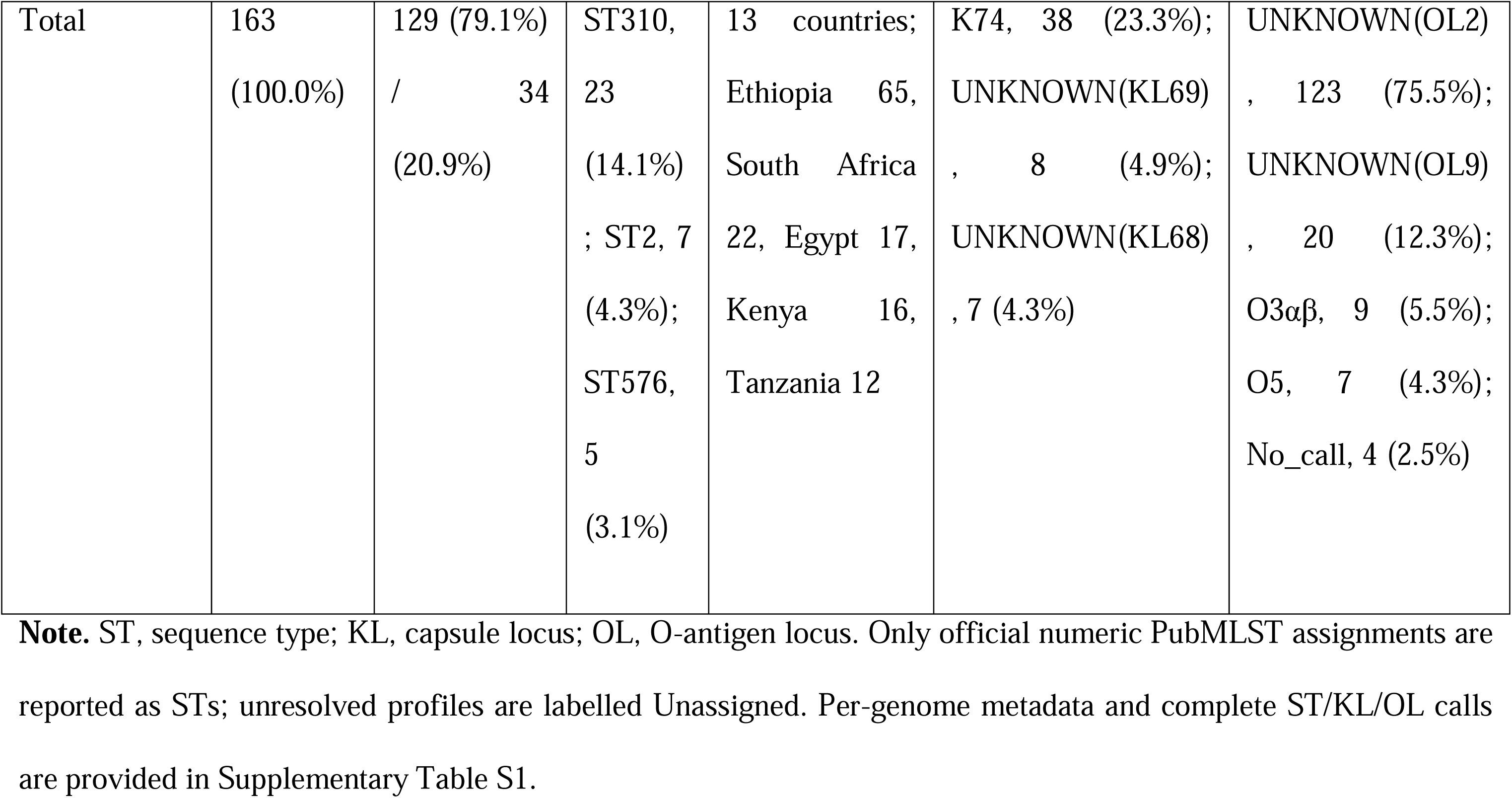
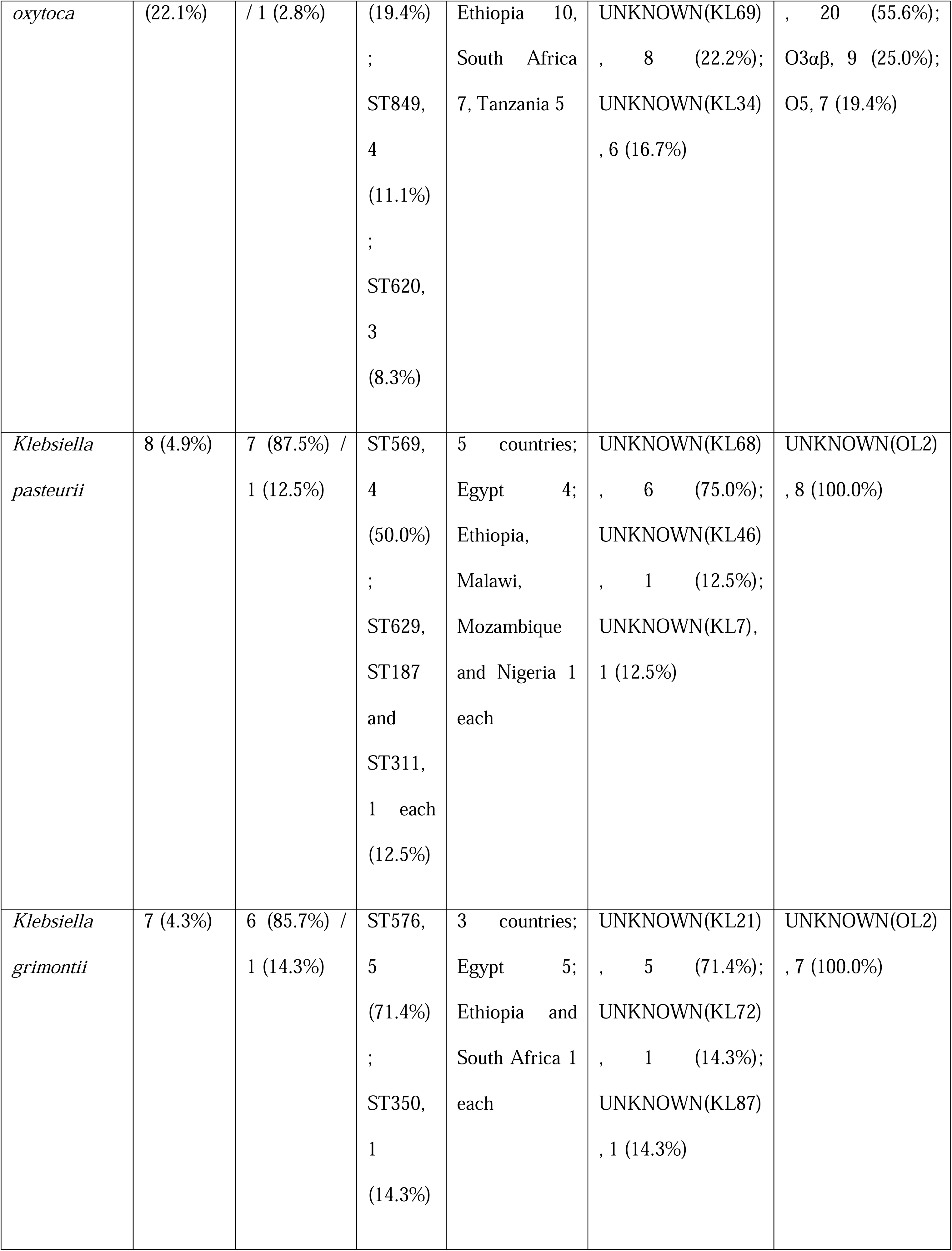
Species composition, current official PubMLST sequence-type status and Kaptive surface-locus calls among 163 African KoSC genomes.

Forty-six KL categories occurred across the African set, with K74 the most frequent call (38/163, 23.3%). O-locus calls were more concentrated: UNKNOWN(OL2) occurred in 123/163 genomes (75.5%), UNKNOWN(OL9) in 20/163 (12.3%), O3αβ in 9/163 (5.5%), O5 in 7/163 (4.3%), and no-calls in 4/163 (2.5%) (Table 1; Supplementary Table S1).

### Species distribution of *til* and *leup* BGCs

Intact *til* BGCs occurred in 55/163 genomes (33.7%; Fig. 3). Four additional genomes contained assembly-fragmented candidates, three disrupted candidates and two unresolved candidates, while 99 had no qualifying *til* locus (Supplementary Table S6). Intact *til* BGCs occurred in all seven *K. grimontii* genomes, 26/36 *K. oxytoca*, 7/8 *K. pasteurii* and 15/112 *K. michiganensis*.

Intact *leup* BGCs occurred in 103/163 genomes (63.2%; Fig. 3; Supplementary Tables S7 and S8), comprising 102/112 *K. michiganensis* and 1/8 *K. pasteurii*. Four additional *K. michiganensis* genomes contained assembly-fragmented candidates, while 56 genomes had no qualifying *leup* locus, including all 36 *K. oxytoca* and all seven *K. grimontii* genomes. Species-by-country strata retained the same broad pattern: for example, 49/53 Ethiopian *K. michiganensis* genomes carried intact *leup*, while 7/10 Ethiopian *K. oxytoca* genomes carried intact *til* (Supplementary Table S5).

### AMR-associated determinants and mobilome features

The AMR matrix contained 54 displayed AMR-associated features (Fig. 4; Supplementary Tables S9–S12). The four core/intrinsic loci *ompA*, *oqxA*, *oqxB* and *bla*_OXY_ occurred in all 163 genomes. Among acquired determinants, *dfrA* was most frequent (101/163, 62.0%), followed by *bla*_CTX-M_ (77/163, 47.2%), *aph(3*″*)-Ib* and *aph*(*6*)*-Id* (75/163 each, 46.0%), *bla*_TEM_ (74/163, 45.4%), *aac(6*′*)-Ib* (70/163, 42.9%), *sul2* (69/163, 42.3%), *aac*(*3*)*-II* (66/163, 40.5%) and *bla*_OXA_ (63/163, 38.7%). *qnrB* and *catB3* each occurred in 57/163 genomes (35.0%), *tet(A)* in 56/163 (34.4%), and *qacE*Δ*1* in 46/163 (28.2%). Carbapenemase-associated determinants included *bla*_NDM_ in 9/163 genomes (5.5%) and *bla*_KPC-2_ and *bla*_VIM-1_ in 1/163 genome each (0.6%) (Supplementary Table S17).

**Figure 4.**
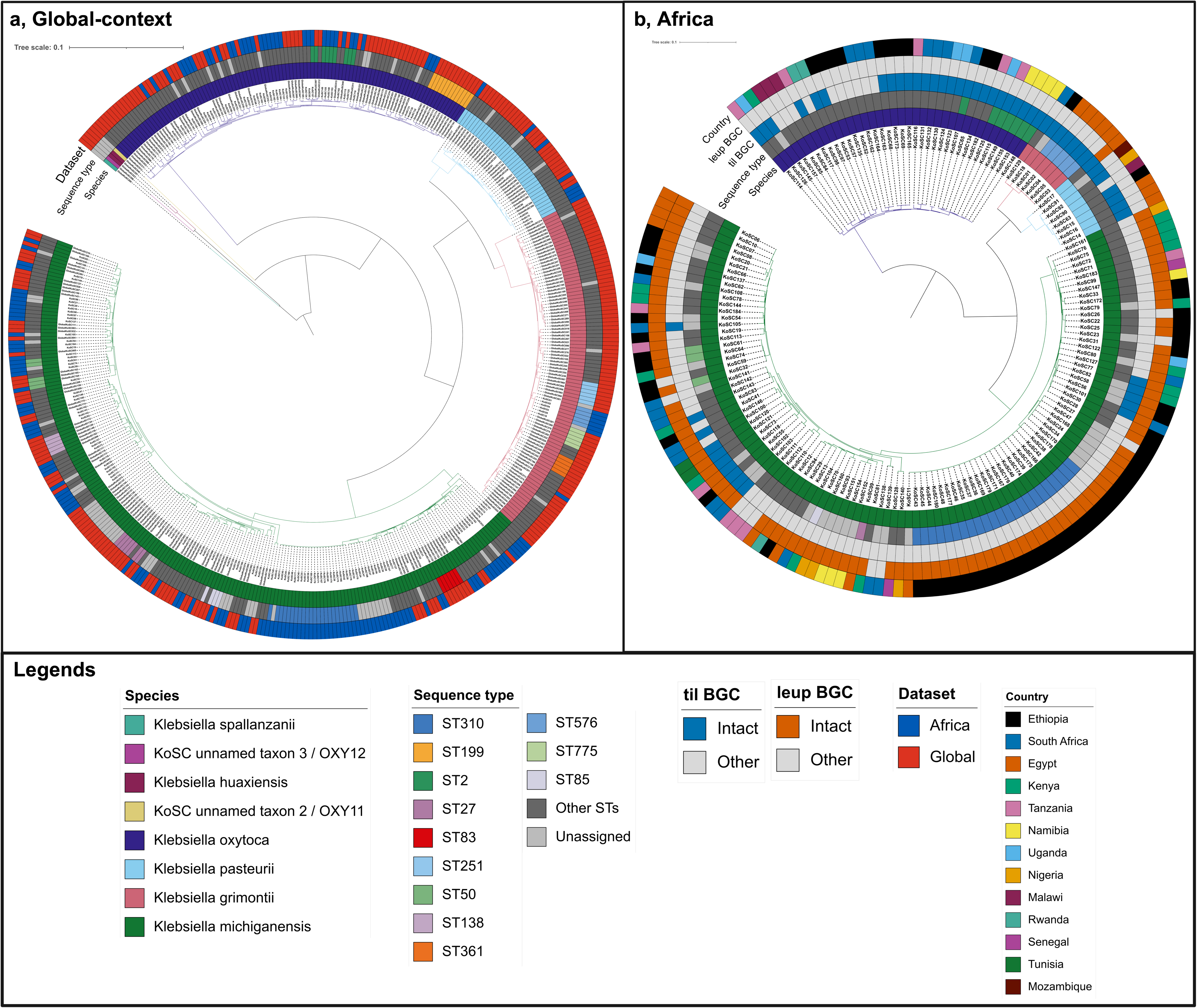
AMR-associated loci, plasmid replicons and non-plasmid MGEs across 163 African KoSC genomes. The persistent/core-gene phylogeny is aligned with species, current official ST, K-locus and O-locus metadata and binary feature matrices. The AMR block contains four core/intrinsic AMR-associated loci and 50 non-core features grouped by antimicrobial class; plasmid-replicon and non-plasmid MGE blocks contain 17 displayed features each. Filled cells indicate sequence detection under the feature-specific criteria described in Methods; yellow cells denote plasmid-replicon detections and pink cells denote non-plasmid MGE detections. Feature prevalence and nomenclature are provided in Supplementary Tables S9–S12 and S17.

The most frequently displayed plasmid replicons were Col(pHAD28) (73/163, 44.8%), IncFIB(K) (64/163, 39.3%), repB(R1701) (36/163, 22.1%), IncFII(Yp) (33/163, 20.2%) and IncFII(p14) (26/163, 16.0%). Frequent non-plasmid MGEs included ISKpn26 (72/163, 44.2%), ISEcp1 and ISKpn34 (69/163 each, 42.3%), ISKpn21 (60/163, 36.8%), ISEcl1 (59/163, 36.2%) and IS903 (55/163, 33.7%) (Fig. 4; Supplementary Table S17).

### VFDB component landscape

*Klebsiella*-focused VFDB screening detected at least one curated reference component in 112/163 genomes. Thirteen of 122 unique curated components were detected across five of 14 factor groups (Supplementary Tables S13–S15). The most frequent component-level signals were *allR* (70/163, 42.9%), *entB* (51/163, 31.3%), *fyuA* (38/163, 23.3%), *ybtA* (32/163, 19.6%), *allB* (27/163, 16.6%), and *allC* (14/163, 8.6%). No genome recovered every curated VFDB component for any of the 14 factor groups.

Targeted type 3 fimbrial architecture analysis identified complete *mrkABCDF* structural-operon candidates in KoSC147, KoSC41 and KoSC55: all *K. michiganensis*. KoSC102 and KoSC103 contained partial *mrkCDF* structural clusters lacking *mrkA* and *mrkB* (Supplementary Table S16).

## DISCUSSION

The African KoSC collection revealed substantial genomic diversity but also a pronounced imbalance in the public sequence record, with *K. michiganensis* dominating the dataset and a large proportion of genomes originating from a small number of countries (Fig. 3; Table 1). This uneven representation is important for interpreting all downstream distributions. Recent continental genomic-surveillance programmes have substantially increased bacterial whole-genome sequencing in Africa but continue to report gaps in metadata completeness and uneven sequencing capacity, while a recent analysis of publicly available African *K. pneumoniae* species-complex genomes similarly found strong country-level sampling imbalance (7, 27). The country and species frequencies observed here therefore define the composition of the available African KoSC genomic record rather than population-level prevalence.

Gene-content diversity was a major feature of the African KoSC population. The predominance of shell and cloud families and their further expansion after inclusion of the global comparator genomes indicate substantial variation outside the conserved gene repertoire (Fig. 2; Supplementary Figure S2). This is consistent with earlier KoSC genomic analyses: Moradigaravand et al. (2017) identified a large accessory genome in *K. oxytoca* (28), whereas Cosic et al. (2021) demonstrated that accessory-gene content varies across KoSC species and is strongly associated with core-genome phylogeny (29). The present persistent/core-gene phylogeny similarly separated the African genomes into species-associated backgrounds within the broader global diversity (Fig. 3). Although most genomes had an official PubMLST assignment, 31/112 *K. michiganensis* genomes remained unassigned (Table 1), indicating that a substantial fraction of the African *K. michiganensis* collection is not represented by current official seven-locus ST definitions.

The African resistome combined conserved KoSC-associated loci with a heterogeneous repertoire of acquired resistance determinants (Fig. 4). The ubiquity of *bla*_OXY_ and *oqxAB* is consistent with their established status as intrinsic KoSC resistance determinants, although *oqxAB* is not conserved across every species currently assigned to the wider complex (3). Acquired β-lactam, aminoglycoside, trimethoprim, sulfonamide, quinolone and tetracycline resistance determinants were distributed across the African genomes, while carbapenemase genes were confined to a smaller subset. This spectrum is concordant with previous KoSC genomic and clinical literature documenting acquired *bla*_CTX-M_, *bla*_TEM_, *bla*_SHV_, *bla*_NDM_, *bla*_KPC_, *bla*_VIM_ and other resistance families in addition to chromosomal *bla*_OXY_ (3, 30). Because antimicrobial susceptibility data were not available for these genomes, the analysis establishes predicted resistance-gene carriage.

The recurrent plasmid replicons and non-plasmid MGEs provide additional context for this acquired resistance diversity (Fig. 4). Previous KoSC studies provide direct evidence that resistance determinants can move within shared plasmid backgrounds: Stewart et al. (2022) identified the same *bla*_IMP-4_-carrying IncL/M plasmid in *K. michiganensis* and *K. pasteurii* (31), while Moradigaravand et al. (2017) found evidence of exchange of accessory resistance and virulence genes between *K. oxytoca* and *K. pneumoniae* (28). The replicon and MGE distributions observed here identify genomic backgrounds in which such mobile elements occur, but they do not by themselves demonstrate plasmid transmission or assign individual resistance genes to complete plasmids. This distinction is particularly important for draft genomes because repetitive plasmid sequences complicate reconstruction from short reads (23).

The contrasting distribution of the *til* and *leup* loci was the strongest species-associated biosynthetic pattern in the African collection (Fig. 3). The intact *til* BGC was concentrated in *K. oxytoca*, *K. pasteurii* and *K. grimontii*, whereas the intact *leup* BGC occurred almost exclusively in *K. michiganensis*. Independent genomic surveys support both components of this pattern. Shibu et al. (2021) identified the complete *til* (kleboxymycin) BGC in *K. oxytoca*, *K. pasteurii*, *K. grimontii* and *K. michiganensis* (8), while McCartney and Hoyles (2025) found the complete *leup* operon in 1,176/1,298 (90.6%) *K. michiganensis* genomes they screened and reported an almost exclusive association of the locus with that species (1). The biological distinction between these loci is supported experimentally. Both tilimycin and tilivalline are cytotoxic pyrrolobenzodiazepine non-ribosomal peptides that contribute to the aetiology of antibiotic-associated haemorrhagic colitis. Tilimycin is genotoxic, whereas tilivalline – produced from tilimycin in the presence of indole – binds tubulin and stabilizes microtubules (9). The *leup* pathway encodes leupeptin biosynthesis and was enriched among respiratory-associated *Klebsiella* isolates in the original pathway study (10); subsequent work showed that *N*-acetylneuraminic acid activates the *leup* operon and production of leupeptin and a derived pyrazinone in *K. michiganensis* (11). Leupeptin produced by *K. michiganensis* is thought to increase survival of the bacterium during infections because leupeptin inhibits a subunit of the immunoproteasome, thereby decreasing immune activity (11). The present analysis establishes genomic locus architecture and distribution of these virulence-associated BGCs in an African context.

The VFDB analysis requires particularly careful interpretation. Under the curated component and factor definitions applied here, only a restricted subset of *Klebsiella*-labelled VFDB components was recovered, and no genome satisfied all components of a complete curated factor (Supplementary Tables S13–S15). This result is not directly equivalent to previous broad virulence-gene screens. Long et al. (2022), using a different reference panel of 80 *K. pneumoniae*-associated virulence genes, detected 72 genes among 588 KoSC genomes and reported individual *mrkABCDFHJ* components in almost all genomes (30). In the present study, the separate *mrk* analysis required *mrkABCDF* to occur together on one contig with consistent order and orientation; only three genomes met that complete structural criterion, with two additional partial clusters (Supplementary Table S16). Thus, the three complete candidates represent stringent recovery of the defined structural operon and do not imply that type 3 fimbrial homologues are absent from the remaining genomes. It should be noted that the VFDB core dataset largely comprises data for experimentally validated virulence factors associated with *K. pneumoniae*. While belonging to the same genus, it is possible that the KoSC may encode virulence factors distinct from other *Klebsiella* species. Support for this statement comes from the exclusive association of the *til* and *leup* virulence-associated BGCs with the KoSC.

Three properties of the dataset delimit the conclusions that can be drawn from these analyses. First, the geographic composition of publicly available African KoSC genomes is highly uneven, precluding population-level prevalence estimates. Second, draft short-read assemblies limit complete reconstruction of plasmids and fragmented genomic loci. Third, antimicrobial-susceptibility and metabolite-production measurements were not linked to the genomes, so resistance-gene detection and BGC architecture cannot be translated directly into phenotypic resistance or metabolite production. Expanded, systematically sampled African collections combining complete metadata, long-read or hybrid assemblies and linked phenotypic measurements will enable these genomic patterns to be tested at population and functional levels.

## CONCLUSION

The African KoSC public-genome collection contains broad species, lineage and accessory-gene diversity, with frequent acquired AMR determinants and heterogeneous plasmid-replicon and MGE profiles. Intact *til* and *leup* BGCs show distinct species distributions, and *Klebsiella*-focused VFDB screening identifies a limited set of additional component signals, including three complete *mrkABCDF* structural-operon candidates. The curated 163-genome African dataset, 282-genome global context panel and linked supplementary audits provide a reproducible genomic baseline for expanded sampling, closed-genome reconstruction and phenotype-linked KoSC studies in Africa.

## DATA AVAILABILITY

This study analysed publicly available sequence data. Accession-level metadata for the African and global-context genomes are provided in Supplementary Table S1. Quality-control, taxonomic, typing, BGC, AMR, mobilome and VFDB results are supplied in Supplementary Tables S2–S17 and Supplementary Figures S1–S3. No new clinical specimens, participant recruitment or animal experiments were included in this study.

## CRediT authorship contribution

**S.Y.B:** Conceptualization, Methodology, Software, Validation, Formal analysis, Investigation, Data curation, Visualization, Writing – original draft, Writing – review & editing. **A. M:** Methodology, Validation, Writing – review & editing. **L. H.:** Methodology, Data curation, Supervision, Validation, Writing – review & editing. **R. S. M.:** Conceptualization, Methodology, Supervision, Project administration, Resources, Writing – review & editing.

