## Supplementary Figure S1 to S3 for "Genome-resolved surveillance of African *Klebsiella oxytoca* species complex genomes reveals resistome-mobilome and biosynthetic gene cluster diversity"

### Supplementary Figures

#### Supplementary Figure S1.

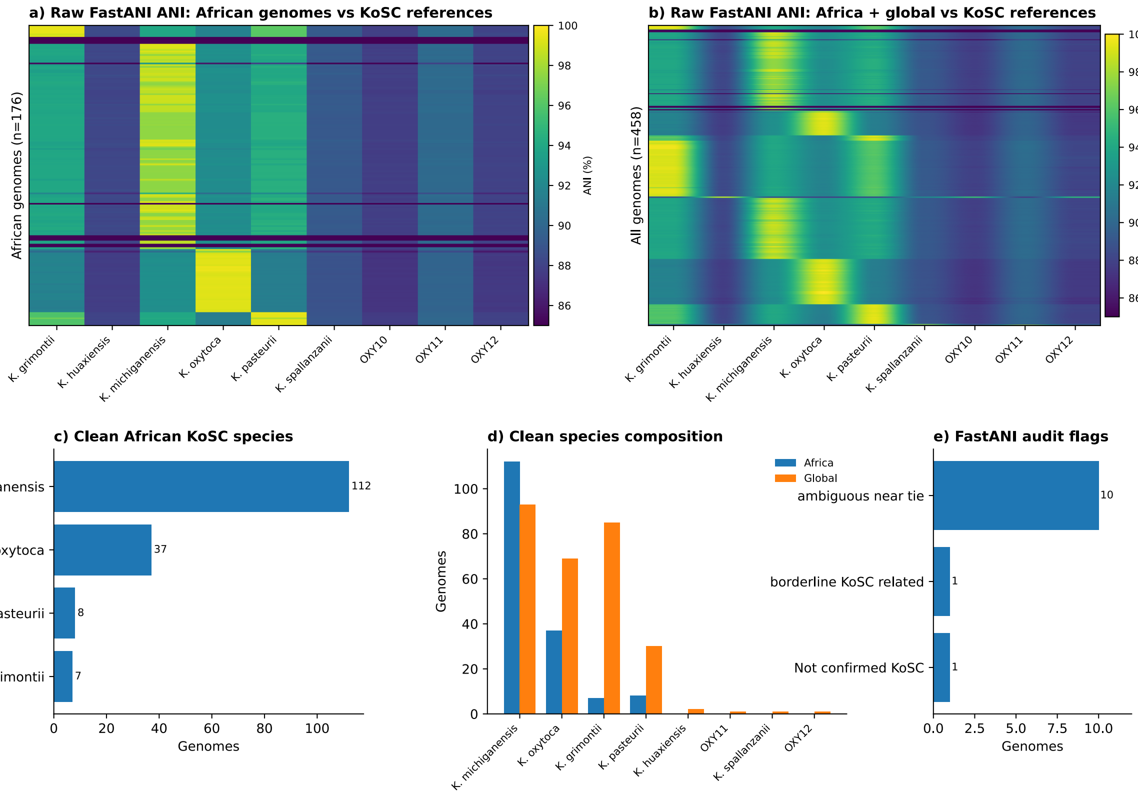


**Figure S1. FastANI-based taxonomic confirmation and audit of the African and global *Klebsiella oxytoca* species complex collections.** The panels show the raw African and combined ANI matrices, taxonomically cleaned African species counts, species composition by dataset, and counts of records flagged during taxonomic reconciliation. The taxonomy-confirmed collection comprised 164 African and 282 global genomes; annotation-dependent African analyses subsequently used 163 genomes after exclusion of KoSC109.

#### Supplementary Figure S2.

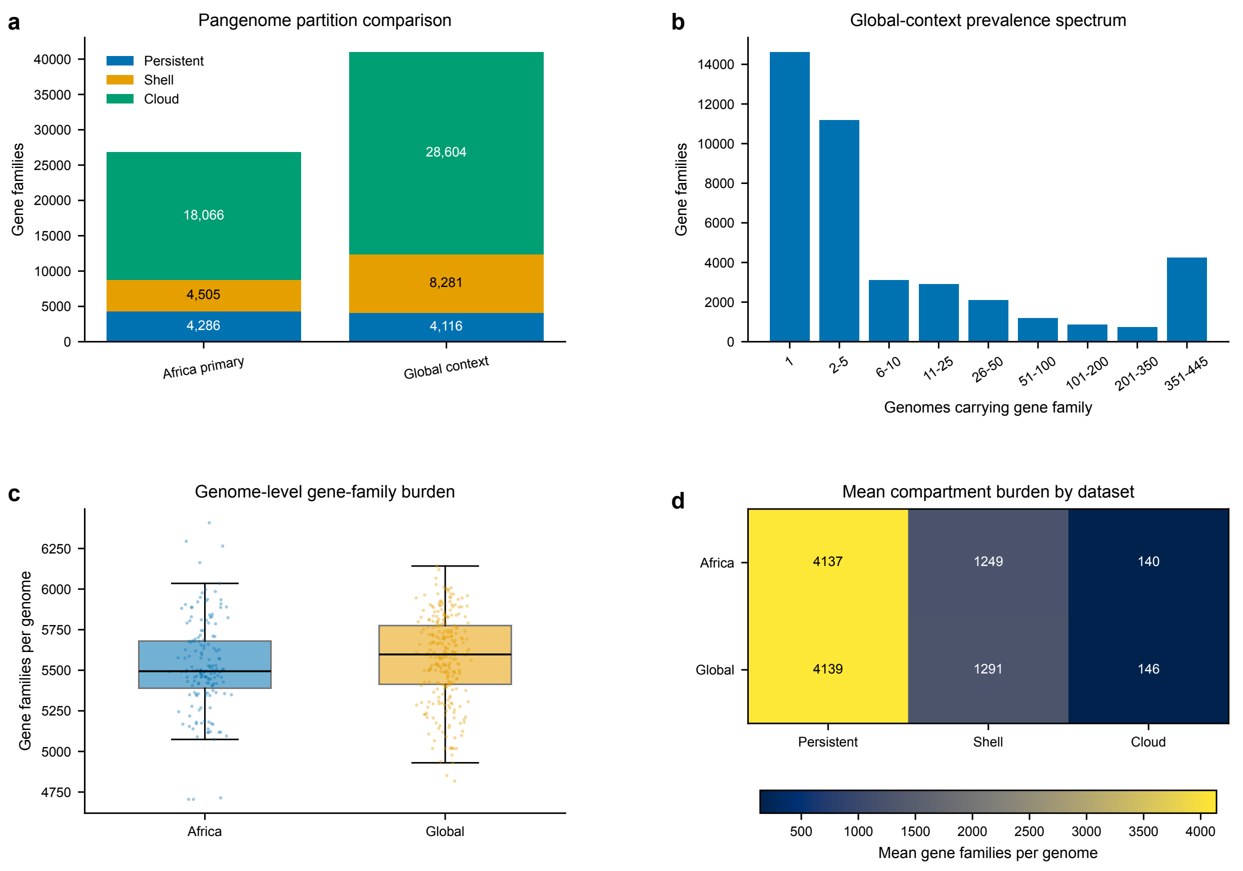


**Figure S2. Global-context pangenome structure of the Klebsiella oxytoca species complex**. Panels compare the African 163-genome pangenome with the combined 445-genome Africa-plus-global analysis, show the combined gene-family frequency spectrum, summarize genome-level gene-family burden by dataset, and report mean persistent, shell, and cloud family burden. The global-context analysis used assembly-derived PPanGGOLiN outputs and excluded KoSC109.

#### Supplementary Figure S3

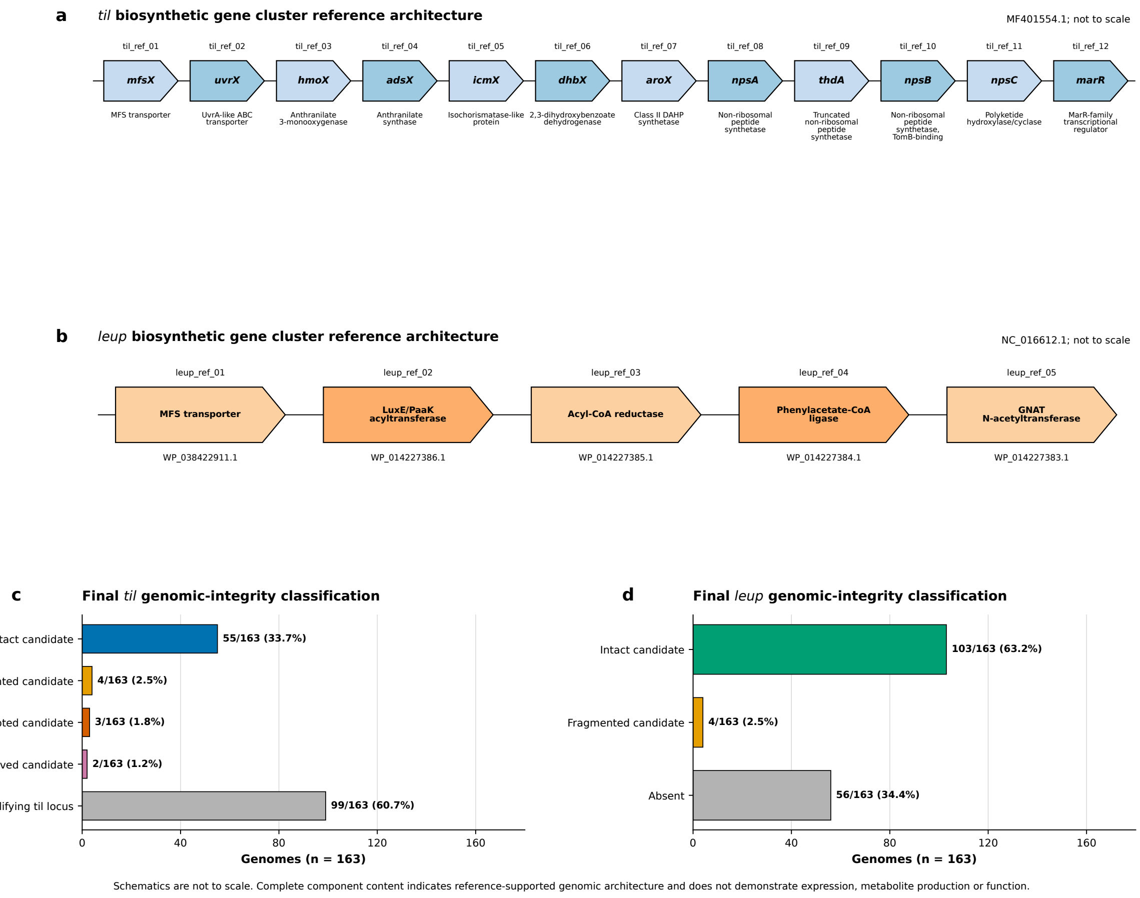


**Figure S3. Reference architecture and genomic-integrity classification of the *til* and *leup* biosynthetic gene clusters in the 163-genome African annotation set.** The figure presents the 12-component *til* and five-component leup reference architectures and the final locus-integrity distributions. The *til* classifications comprised 55 intact, four assembly-fragmented, three disrupted, two unresolved, and 99 genomes without a qualifying locus. The *leup* classifications comprised 103 intact, four fragmented, and 56 absent loci. Intact genomic architecture does not establish transcription, metabolite production or biological activity.
